# Actomyosin-mediated cellular tension promotes Yap nuclear translocation and myocardial proliferation through α5 integrin signaling

**DOI:** 10.1101/2022.06.09.495549

**Authors:** Xiaofei Li, Callie McLain, Michael S. Samuel, Michael F. Olson, Glenn L. Radice

## Abstract

The cardiomyocyte phenotypic switch from a proliferative to terminally differentiated state results in the loss of regenerative potential of the mammalian heart shortly after birth. Yet, the molecular mechanisms that regulate this critical developmental process are incompletely understood. Nonmuscle myosin IIB (NM IIB)-mediated actomyosin contractility regulates cardiomyocyte cytokinesis in the embryonic heart, and NM IIB levels decline after birth suggesting a role for cellular tension in the regulation of cardiomyocyte cell cycle activity in the postnatal heart. The Rho kinase (ROCK) serine/threonine protein kinases that act downstream of the RhoA small GTP-binding protein regulate nonmuscle myosin contractile force generation. To investigate the role of actomyosin contractility in cardiomyocyte maturation and cell cycle arrest, we conditionally-activated ROCK2 kinase domain (ROCK2:ER) in the murine postnatal heart. Here we show that cardiac-specific activation of actomyosin contractility shifts the balance from cell-cell to cell-matrix adhesions. Specifically, α5/β1 integrin and fibronectin matrix increase in response to actomyosin-mediated tension. Moreover, activation of ROCK2:ER promotes nuclear translocation of Yap, a mechanosensitive transcriptional co-activator, and enhances cardiomyocyte proliferation. Finally, we show that reduction of myocardial α5 integrin rescues the myocardial proliferation phenotype in ROCK2:ER hearts. These data demonstrate that cardiomyocytes respond to increase intracellular tension by altering their intercellular contacts in favor of cell-matrix interactions leading to Yap nuclear translocation, thus uncovering a novel function for nonmuscle myosin contractility in promoting cardiomyocyte cell cycle activity in the postnatal heart.

## Introduction

The neonatal mammalian heart is capable of substantial regeneration following injury; however this regenerative capacity largely disappears in the first week after birth (Porrello et al., 2011; Porrello et al., 2013). As such, the adult human heart does not have an effective mechanism to regenerate the cardiomyocytes (CMs) lost following a heart attack. Instead, the lost muscle is replaced by the excessive deposition of extracellular matrix (ECM), which together reduces both systolic and diastolic left ventricular (LV) functions of the heart and often leads to heart failure. Thus, a better understanding of the molecular mechanisms controlling CM proliferation is required in order to identify novel targets that will stimulate cardiac regeneration following myocardial infarction.

The Rho-associated coiled-coiled containing protein serine/threonine kinases ROCK1 and ROCK2 are downstream effectors of the Rho subfamily of small GTPases (Hartmann et al., 2015; Noma et al., 2006; Shimokawa et al., 2016). As a key regulator of intracellular contractility, ROCK allows cells to respond to mechanical cues from the tissue microenvironment. Evidence from animal studies indicate an important role for ROCK in cardiac morphogenesis and in adult heart pathophysiology (Shi et al., 2011). Pharmacological inhibition of ROCK during early chick and mouse embryogenesis caused defects in cardiac morphogenesis (Wei et al., 2001). Consistent with this finding, simultaneous deletion of both ROCK1 and ROCK2 in the heart using cardiac-specific αMHC-Cre transgene caused embryonic lethality (Shi et al., 2019). Treatment of adult rats with the ROCK inhibitor, fasudil, blocks angiotensin II-induced cardiovascular hypertrophy (Higashi et al., 2003). Moreover, fasudil suppressed cardiomyocyte hypertrophy and interstitial fibrosis after myocardial infarction in mice (Hattori et al., 2004). Global knockout of ROCK2 in mice results in placental dysfunction, intrauterine growth retardation, and fetal death (Thumkeo et al., 2003). Interestingly, cardiac-restricted knockout of ROCK2 using αMHC-Cre transgene (ROCK2 CKO mouse model) did not result in a cardiac phenotype under basal conditions (Okamoto et al., 2013; Sunamura et al., 2018). However, following angiotensin II treatment or transverse aortic constriction (TAC) ROCK2 CKO mice exhibited less cardiac hypertrophy, reduced end-diastolic wall thickness, and decreased left ventricular (LV) mass compared to control littermates supporting the idea that ROCK2 is critical for CMs to respond to mechanical stimuli. Using a ROCK dominant-negative transgenic model (ROCK DN), endogenous ROCK activity was blocked in embryonic CMs resulting in disruption of sarcomeres and reduced CM proliferation (Bailey et al., 2019), although no molecular mechanism was provided to account for the decreased CM proliferation. These *in vivo* studies demonstrate the requirement for ROCK2 signaling in cardiac development and cardiac remodeling following mechanical stress, but its specific role in postnatal CM maturation remains poorly defined.

ROCK mediates the phosphorylation of myosin regulatory light chains leading to increased myosin ATPase activity, thus stimulating actomyosin contractility. Nonmuscle myosin IIB (NM IIB) is the primary NM isoform expressed in heart muscle. As CMs mature after birth, nonmuscle myosin activity decreases and it is reactivated in response to mechanical stress in the adult heart, e.g., after myocardial infarction (Ma and Adelstein, 2012; Pandey et al., 2018). Germline knockout of NM IIB in mice leads to multiple developmental abnormalities including cardiac malformations and cardiac hypertrophy resulting in embryonic lethality (Tullio et al., 1997). Notably, defective cytokinesis causes an increased number of enlarged binucleated CMs in the NM IIB mutant hearts, supporting a role for NM IIB in myocardial proliferation (Takeda et al., 2003). Mechanistically, NM IIB actin cross-linking activity is required to generate tension that drives cytokinesis (Ma et al., 2012). Cardiac-specific NM IIB knockout (αMHC-Cre) mice also exhibit cytokinesis defects; however, the mutant mice survive and go on to develop progressive cardiomyopathy with age (Ma et al., 2009). Taken together, these data indicate the importance of NM IIB in regulating cardiac development and homeostasis including CM cytokinesis.

Mechanical equilibrium between cells and tissues is dependent on maintaining a proper balance between cell-cell and cell-ECM interactions (Han and de Rooij, 2016; Mui et al., 2016; Zuidema et al., 2020). This is especially evident in the heart where the contractile force generated by myocytes provides mechanical stimuli both to the individual myocyte and to neighboring cells through mechanosensing structures, including cadherin- and integrin-based adhesion complexes. Despite the importance of these two mechanosensing structures to the function of cells and organs, little is known regarding the reciprocity between integrin adhesions and cadherin adhesions in the regulation of cardiac development and homeostasis.

The transition from hyperplastic to hypertrophic growth in the postnatal heart is accompanied by dynamic remodeling of cell-cell and cell-ECM adhesive junctions. Specifically, CM α5/β1 integrin-fibronectin adhesions decrease after birth while N-cadherin junctions remodel and accumulate at myocyte termini leading to the formation of a specialized junction called the intercalated disc (ICD) (Vite and Radice, 2014). Establishment of functional cadherin and integrin adhesion complexes require the reorganization and assembly of the actin cytoskeletal network. Modulation of intracellular tension can have different effects on cell-cell and cell-ECM interactions depending on the cellular context (de Rooij et al., 2005; Dzamba et al., 2009; Liu et al., 2010; Martinez-Rico et al., 2010; Shewan et al., 2005). Here, we focus on the effects of cellular tension on the remodeling of myocardial adhesions in the postnatal heart prior to ICD assembly.

We previously reported that disrupting the linkage between N-cadherin and the actin cytoskeleton by depleting αE-/αT-catenins caused an increase in RhoA activity, mislocalization of actomyosin contractility, and enhanced CM proliferation in the postnatal mouse heart (Vite et al., 2018). Whether enhanced intracellular tension is responsible for the CM hyperproliferation phenotype in the αE-/αT-catenin mutant mice is unknown. To address this possibility, here we studied the direct effect of activating ROCK2 signaling and consequent actomyosin contractility on CM maturation. We find that inducing actomyosin contractility leads to a rebalancing of myocardial cell-cell and cell-matrix adhesions, with the latter responsible for Yap nuclear translocation and CM proliferation. These results implicate the actomyosin cytoskeleton as a critical mediator of cross talk between cadherin and integrins, and show that activating nonmuscle myosin contractility leads to enhanced myocardial cell-fibronectin interactions which stimulate CM cell cycle activity in the postnatal heart.

## Results

### ROCK activation increases non-muscle myosin contractility in the postnatal heart

The temporal expression of non-muscle myosin IIB (NM IIB), non-muscle myosin light chain 2 (MLC2), and active phosphorylated MLC2 (pMLC2) was examined in postnatal hearts of wild-type mice (C57BL/6J) by Western **(Fig. 1A)**. NM IIB, which controls tension within the actin cytoskeleton, is activated by phosphorylation of its light chain at Ser19 (pMLC). Both NM IIB levels and MLC2 activity decline after birth coincident with CM differentiation and cell cycle arrest **(Fig. 1A)**. Whereas ROCK2 levels remain constant over the same period. To investigate the effects of actomyosin contractility on postnatal CMs, we used a conditionally activated ROCK2:ER mouse model outlined in **Figure 1B** (Samuel et al., 2016). Fusion of the ROCK2 kinase domain with the estrogen receptor (ER) hormone-binding domain generates a ROCK2:ER chimeric protein that is inactive in the absence of ligand, but which can be conditionally activated *in vivo* by administering the estrogen analogue tamoxifen (Rath et al., 2017; Samuel et al., 2016). ROCK2:ER mice were bred with αMHC/Cre mice to generate a cardiac-restricted ROCK2:ER mouse model. Western analysis confirmed expression of the ROCK2:ER fusion protein in ROCK2:ER; αMHC-Cre hearts **(Fig. 1C)**. CM proliferation decreases after postnatal day 4 (P4) in mice (Soonpaa et al., 1996). Therefore, to investigate the effects of cellular tension on the transition from hyperplastic to hypertrophic CM growth, mice were administered tamoxifen via IP injection for three consecutive days: P5, P6, and P7. On P7, 4 hrs after the last tamoxifen injection, hearts were harvested for analysis. ROCK2 kinase is required for RhoA-initiated actomyosin contractility through phosphorylation of substrates including MLC2 and MYPT1. Western analysis confirmed increased expression of NM IIB, pMLC2 (Ser19), and pMYPT (Thr696) in ROCK2:ER; MHC-Cre hearts following tamoxifen administration **(Fig. 1D)**. For all subsequent studies, pups were administered tamoxifen as described above, and littermates lacking the Cre transgene served as controls. Actomyosin contractility, as visualized by punctate pMLC immunofluorescence, is higher in P7 ROCK2:ER CMs compared to control **(Fig. 1E)**. Consistent with increased nonmuscle myosin contractility, there is more NM IIB along the lateral membrane of ROCK2:ER CMs compared to control **(Fig. 1F)**.

**Figure 1.**
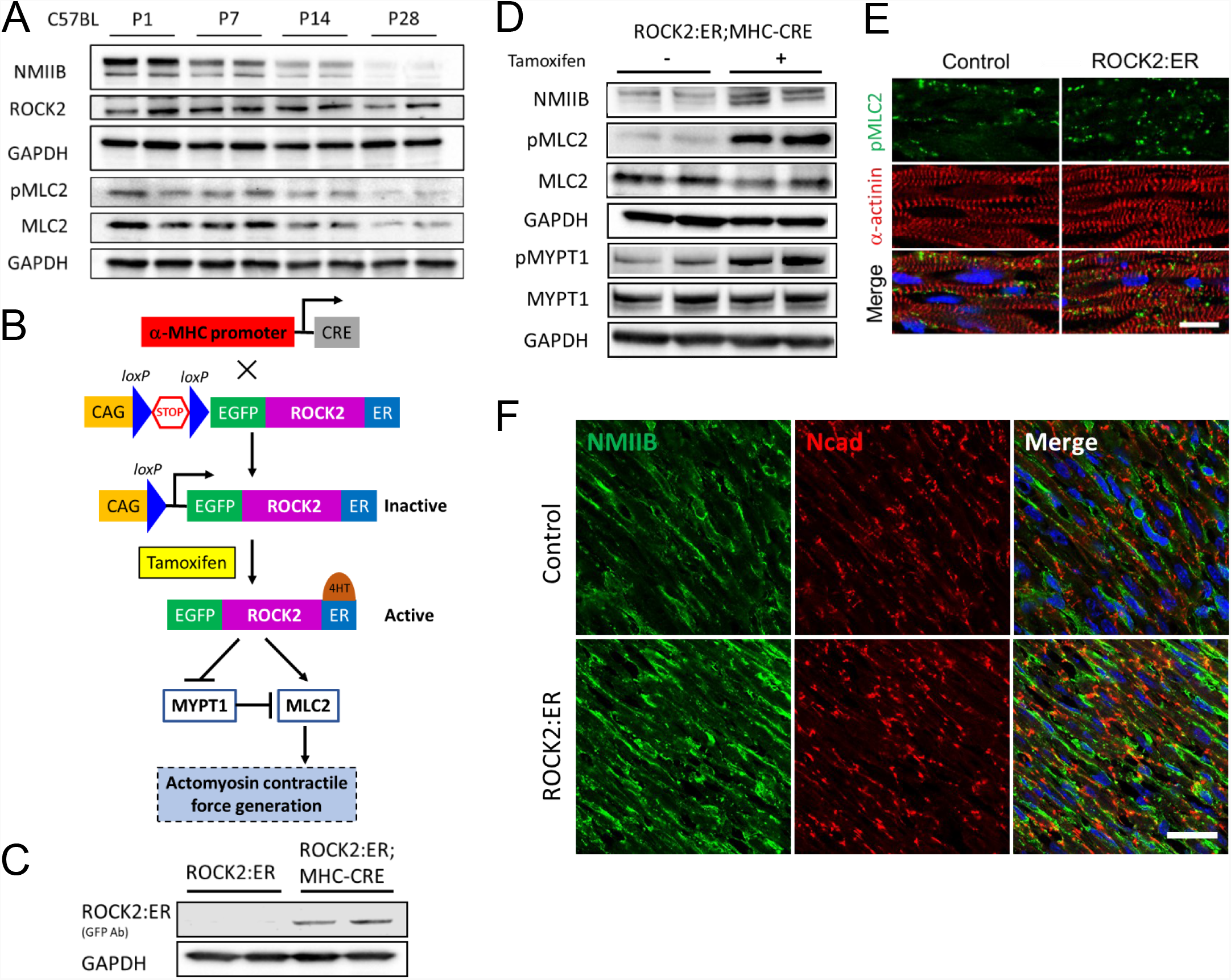
Activation of ROCK2 kinase activity in the postnatal heart. (A) Western blot of NMIIB, ROCK2, pMLC2, MLC2 expression in C57BL wild-type mice at P1, P7, P14, and P28. (B) Schematic of the bitransgenic mouse model consisting of a cardiac-specific Cre transgene and lox-STOP-lox ROCK2:ER transgene. The conditionally-active ROCK2:ER fusion protein is inactive in the absence of ligand. Upon stimulation with 4-hydroxytamoxifen, the specific activity of the kinase domain of ROCK2 increases leading to phosphorylation of various substrates, which results in the generation of actomyosin contractile force. (C) Western blot of ROCK2:ER expression in the ROCK2:ER mouse heart. (D) Western blot analysis of NMIIB, pMLC2, MLC2, pMYPT1, MYPT1 expression in P5 hearts from ROCK2:ER mice before and after tamoxifen activation. (E) Representative immunofluorescent images of heart sections from control and ROCK2:ER mice co-stained with pMLC2 (green) and a-actinin (red). Scale bar: 50μm. (F) Representative immunofluorescent images of heart sections control and ROCK2:ER mice co-stained with NMIIB (green) and N-cadherin (red). Scale bar: 25μm.

### Increased intracellular tension shifts the balance between cell-cell to cell-matrix adhesions

Both the cadherin- and integrin-based adhesion systems are linked to the cortical actin cytoskeleton and share common downstream actin regulatory pathways including the cytoskeletal adaptor protein vinculin. Vinculin regulates cell adhesion by directly binding to actin, stimulating actin polymerization and recruiting actin remodeling proteins in response to intrinsic and extrinsic mechanical forces (Bays and DeMali, 2017). As a mechanosensitive protein, vinculin undergoes a conformational change in response to mechanical force that results in actin remodeling and adhesion strengthening at cell junctions, a prerequisite for cadherin junction maturation. The junction remodeling process requires phosphorylation of vinculin at residue Y822 by the nonreceptor tyrosine kinase c-Abl (Bays et al., 2014). Thus, increased levels of vinculin phosphorylation (Y822) correlate with less mature cadherin junctions. To investigate vinculin activity following ROCK activation, we performed immunofluorescence and Western analyses on P7 hearts using a phospho-specific antibody that recognizes pVCL-Y822 (Bays et al., 2014). Vinculin (VCL) was primarily localized at the membrane as expected, whereas phosphorylated vinculin (pVCL) exhibited a diffuse but heterogeneous distribution, which was enhanced in ROCK hearts compared to control **(Fig. 2A, C)**. Western analysis confirmed the increased pVCL in the ROCK hearts **(Fig. 2B)**. A portion of the pVCL co-localized with N-cadherin at the membrane **(Fig. 2C, arrowheads)**, however most remained diffuse in the cytosol. Total VCL and N-cadherin protein levels did not change in ROCK hearts **(Fig. 2B, D)**. These results are consistent with previous observation that pharmacological inhibition of ROCK blocked force-induced vinculin phosphorylation in cultured epithelial cells (Bays et al., 2014). We previously demonstrated that simultaneous depletion of both αE- and αT-catenin resulted in disruption of nascent N-cadherin junctions, Rho activation, and enhanced actomyosin contractility (Vite et al., 2018). Indeed, we found that pVCL expression was increased in the αE-/αT-catenin double knockout CMs **(Fig. S1)** and comparable to that observed in the ROCK model, providing independent support for pVCL as a marker of dynamic junction remodeling in the postnatal heart. Taken together, these data are consistent with the presence of less mature N-cadherin junctions after activation of ROCK.

**Figure 2.**
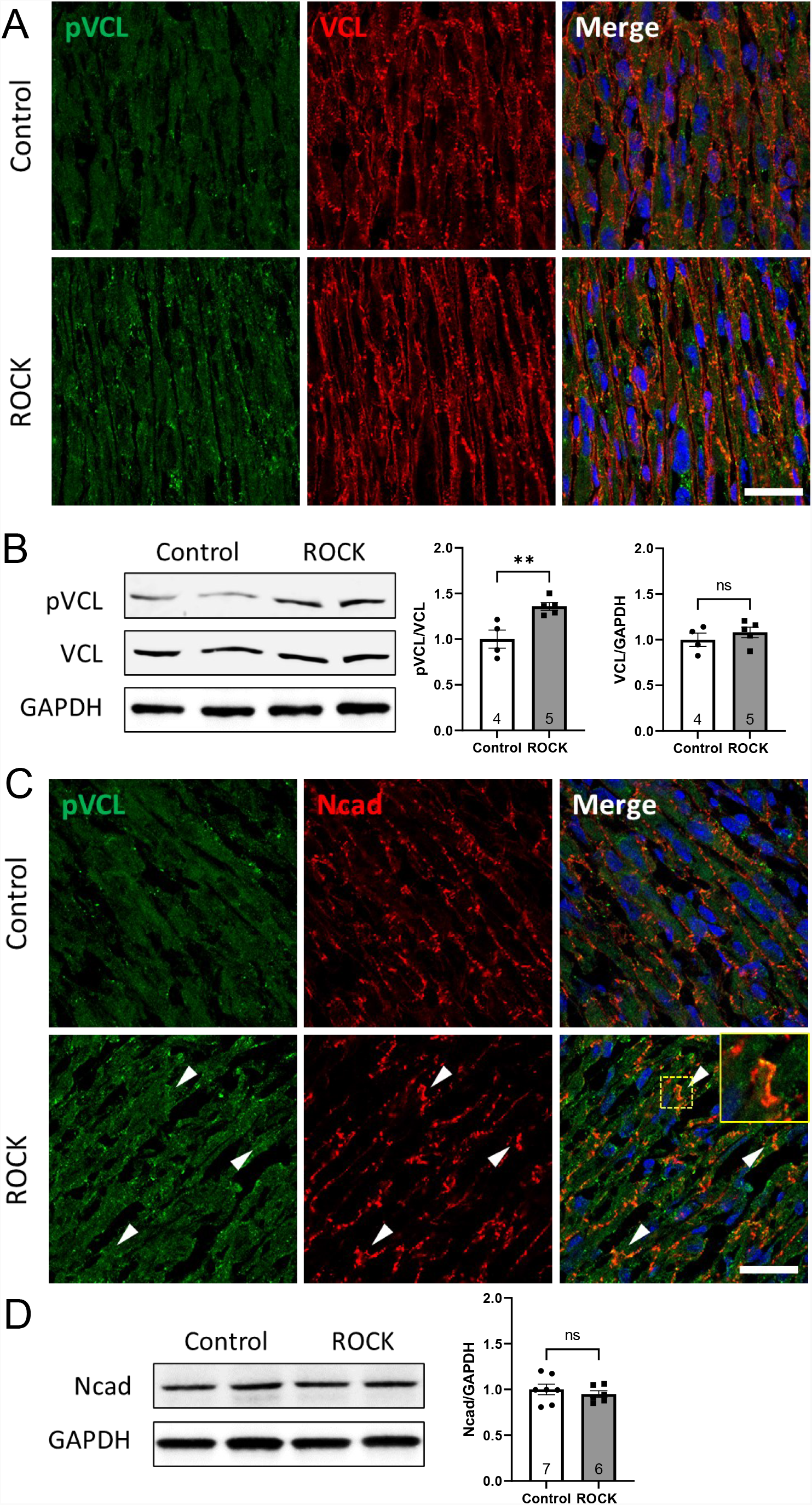
Increased VCL phosphorylation following ROCK activation. (A) Representative immunofluorescence images of heart sections from control and ROCK P7 mice co-stained for pVCL-Y822 (green) and VCL (red). (B) Western blot and quantitative analysis of pVCL and VCL expression in heart tissue lysate from control (n=4) and ROCK (n=5) P7 mice. (C) Representative immunofluorescence images of heart sections from control and ROCK P7 mice co-stained for pVCL-Y822 (green) and N-cadherin (red). pVCL appeared diffuse throughout the cytosol along with its co-localization with N-cadherin at the membrane (arrowheads, inset). (D) Western blot and quantitative analysis of N-cadherin expression in heart tissue lysate from control (n=7) and ROCK (n=6) P7 mice. ns indicates a statistically non-significant difference,** P<0.01 by Student’s t-test. Error bars represent S.E.M.. Scale bar: 25μm.

To investigate the effects of actomyosin contractility on the crosstalk between cell-ECM and cell-cell interactions, we examined the spatial distribution and expression of α5/β1 integrin, the receptor for fibronectin (FN), and N-cadherin in P7 ROCK hearts **(Fig. 3A)**. At this developmental stage, N-cadherin and α5 integrin exhibited extensive co-localization at the membrane with a notable increase in α5 integrin expression at the lateral membrane in the ROCK CMs **(Fig. 3A, enlarged images)**. Western analysis of heart lysates confirmed the increased α5 integrin expression in ROCK hearts **(Fig. 3B)**. The increased α5 integrin expression was accompanied by enhanced extracellular FN matrix along the lateral membrane consistent with strengthening of myocardial cell-matrix interactions in the ROCK hearts **(Fig. 3C)**. Taken together, these data suggest that increased actomyosin contractility shifts the balance from cadherin-based adhesions to integrin-based adhesions in postnatal CMs.

**Figure 3.**
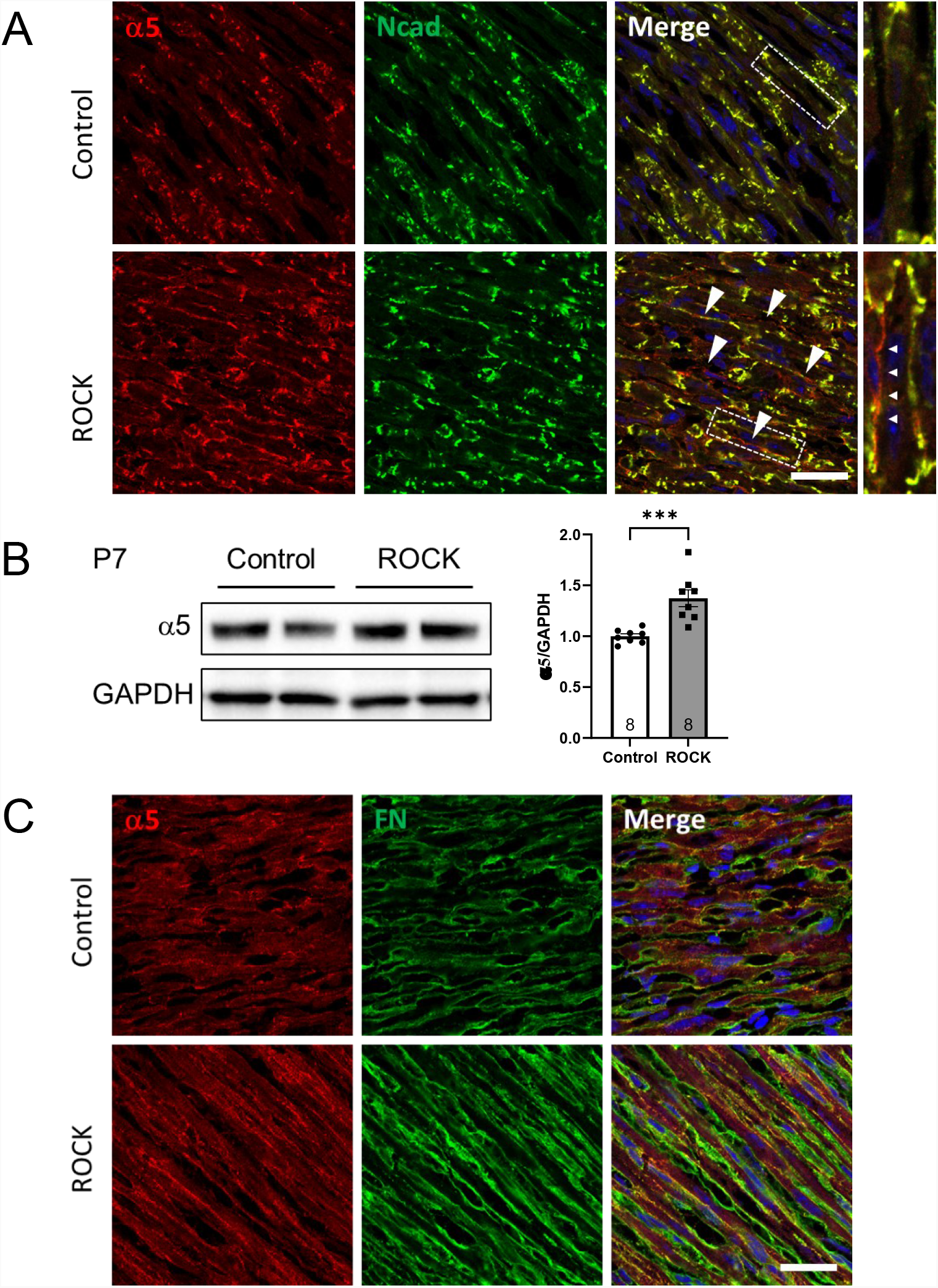
Increased cellular tension leads to enhanced cardiomyocyte α5 integrin-fibronectin interactions. (A) Representative immunofluorescence images of heart sections from control and ROCK P7 mice co-stained for α5 integrin (red) and fibronectin (green). (B) Western blot and quantitative analysis of α5 integrin expression in heart tissue lysate from control (n=8) and ROCK (n=8) P7 mice. (C) Representative immunofluorescence images of heart sections from control and ROCK P7 mice co-stained with α5 integrin (red) and N-cadherin (green). Arrowheads indicate increased lateral α5 integrin expression. Enlarged boxed area shown on right. *** P<0.001 by Student’s t-test. Error bars represent S.E.M.. Scale bars: 25μm.

### ROCK activation drives Yap nuclear translocation and increased cardiomyocyte proliferation

The tissue microenvironment plays a key role in the fate and function of CMs. FN and ECM-associated growth factors are critical regulators of CM proliferation and behavior (Ieda et al., 2009; Wu et al., 2020). Therefore, we examined EdU incorporation (indicator of S phase) and phosphorylation of histone H3 (indicator of M phase) in ROCK hearts at P7 **(Fig. 4A, B)**. The percentage of EdU-positive CMs in ROCK hearts (10.66%±0.70%, n=12) was about 1.5-fold higher than in control (6.90%±0.51%, n=10, *p* < 0.001) P7 hearts. Similarly, the percentage of pHH3-positive CMs in ROCK hearts (8.48%±0.61%, n=11) was about 1.9-fold higher than in control (4.50%±0.24%, n=11, *p* < 0.0001) P7 hearts.

**Figure 4.**
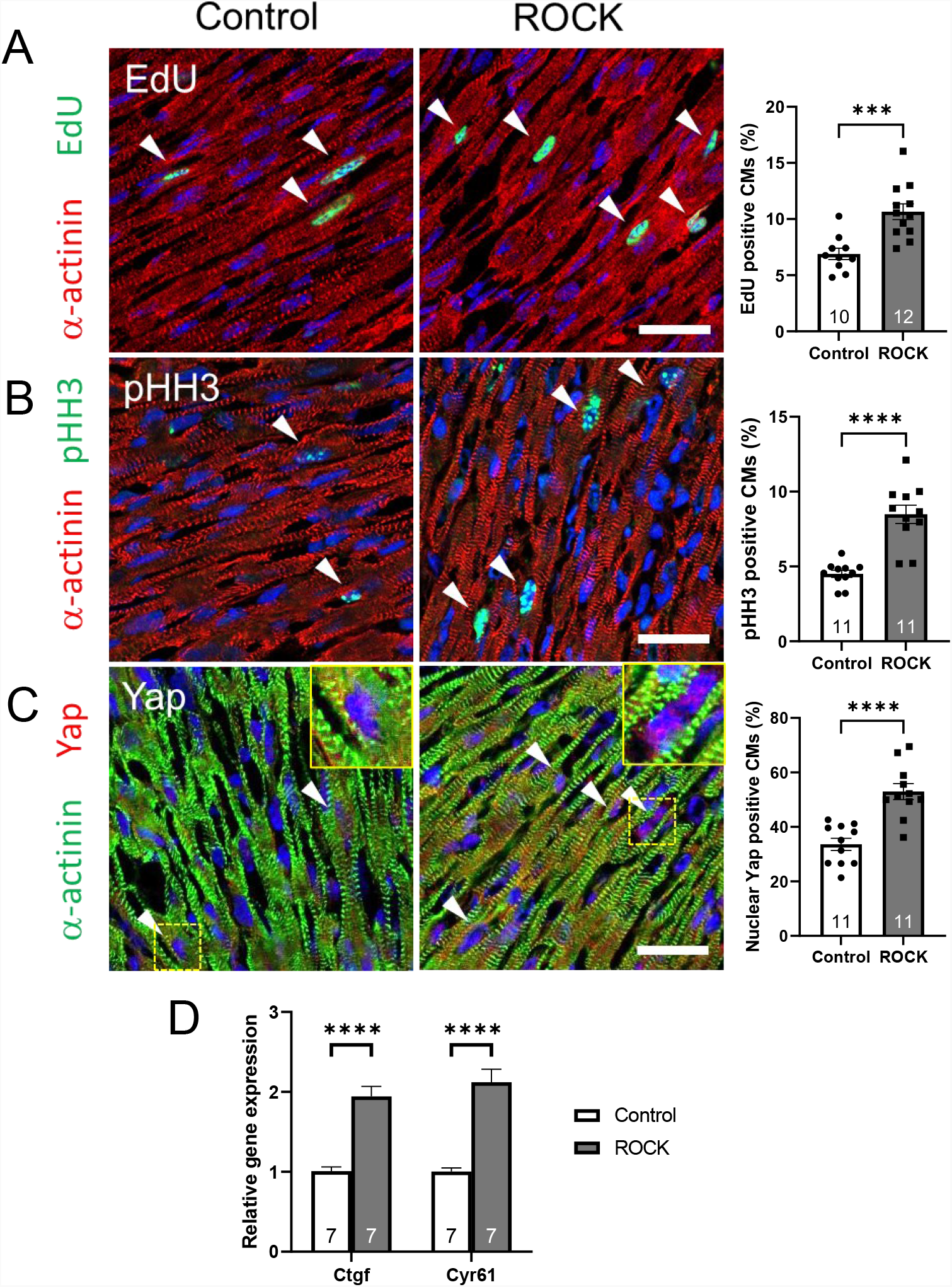
ROCK activation drives Yap nuclear translocation and increased cardiomyocyte cell cycle activity. Representative immunofluorescence images and quantification of heart sections from control and ROCK (n=10-12) P7 mice co-stained with (A) EdU (green)/α-actinin (red), (B) pHH3 (green)/α-actinin (red), and (C) Yap (red)/α-actinin (green). Arrowheads indicate EdU, pHH3, or nuclear Yap-positive cardiomyocytes. (D) Yap target gene expression assessed by quantitative reverse transcription polymerase chain reaction (qRT-PCR) in control (n=7) and ROCK (n=7) P7 hearts. ***, P<0.001; ****, P<0.0001 by Student’s t-test. Error bars represent S.E.M.. Scale bars: 25μm.

Mechanical cues including rearrangement of cytoskeletal tension control Yap activity independently of the Hippo kinase cascade (Dupont et al., 2011), and, importantly, Yap was shown to be necessary and sufficient for CM proliferation (von Gise et al., 2012; Xin et al., 2013). Therefore, we examined the cellular distribution of Yap after activation of actomyosin contractility **(Fig. 4C)**. Quantitative analysis demonstrated that the ROCK hearts had a 1.6-fold increase percentage of nuclear Yap positive CMs (52.97%±2.93%, n=11) compared to control littermates (33.61%±2.23%, n=11, *p*< 0.0001). Next, we examined expression of Yap target genes by quantitative reverse transcription polymerase chain reaction. Yap target genes *Ctgf* and *Cyr61* were upregulated 2-fold consistent with increased Yap transcriptional activity in the ROCK hearts **(Fig. 4D)**. Together, these data indicate that increasing actomyosin cytoskeletal tension is sufficient to drive Yap to the nucleus and increase cell cycle activity supporting a role for nonmuscle myosin contractility in the regulation of CM proliferation in the postnatal heart.

### Cardiomyocyte-fibronectin interactions are required for the proliferation phenotype in ROCK2:ER hearts

ECM and ECM-associated proteins play a critical role in regulating CM proliferation (Ieda et al., 2009; Wu et al., 2020). To determine whether CM-FN interactions are required for the hyperproliferation phenotype in the ROCK hearts, we introduced an *α5 integrin* (*Itga5*) floxed allele into the ROCK2:ER model to generate ROCK2:ER; *Itga5 f/+* (control) and ROCK2:ER; *Itga5 f/+*; MHC-Cre mice **(Fig. 5A)**. Deletion of one copy of *Itga5* resulted in reduced α5 integrin membrane expression in ROCK; α5 +/-CMs **(Fig. 5B)**. Consistent with knockdown of α5 integrin, FN matrix surrounding the ROCK; α5 +/-CMs was reduced **(Fig. 5B)**. Western analysis confirmed reduction of α5 integrin in ROCK; α5 +/-heart lysates **(Fig. 5C)**. Next, we compared CM proliferation rates of ROCK2:ER; *Itga5 f/+* (control), ROCK2:ER; MHC-Cre (ROCK), and ROCK2:ER; *Itga5 f/+*; MHC-Cre (ROCK; α5 +/-) mice **(Fig. 6A, B)**. The percentage of EdU-positive CMs in ROCK hearts (9.25%±0.57, n=7) was significantly higher compared to control (4.82%±0.76%, n=4, *p* < 0.001) and ROCK; α5 +/-hearts (6.78%±0.32%, n=7, *p* < 0.01) at P7. Similarly, the percentage of pHH3-positive CMs in ROCK hearts (7.56%±0.09%, n=6) was increased significantly compared to control (5.31%±0.12%, n=4, *p* < 0.001) and ROCK; α5 +/-hearts (6.01%±0.35%, n=7, *p* < 0.01) at P7. While EdU and pHH3 were not significantly different between ROCK; α5 +/- and control hearts. Thus, deletion of one copy of *Itga5* was sufficient to rescue the excessive mitotic CMs in the ROCK hearts. Next, we assessed Yap cellular distribution in the rescued hearts. Indeed, nuclear translocation of Yap was reduced in ROCK hearts upon deletion of one *Itga5* allele **(Fig. 6C)**. The percentage of nuclear Yap positive CMs in ROCK hearts (56.62%±2.73%, n=6) was increased significantly compared to control (31.23%±2.27%, n=4, *p* < 0.001) and ROCK; α5 +/-CMs (43.49%±3.62%, n=7, *p* < 0.05) at P7. While the percentage of nuclear Yap positive CMs was not significantly different between ROCK; α5 +/- and control hearts. Collectively, our results demonstrate that high nonmuscle myosin contractility shifts the balance from cell-cell to cell-matrix adhesion leading to Yap nuclear translocation and CM proliferation **(Fig. 7)**. Furthermore, genetic knockdown of *Itga5* support the proposition that CM-matrix interactions (i.e., α5/β1 - FN) regulate Yap activity and CM cell cycle activity in the postnatal ROCK heart.

**Figure 5.**
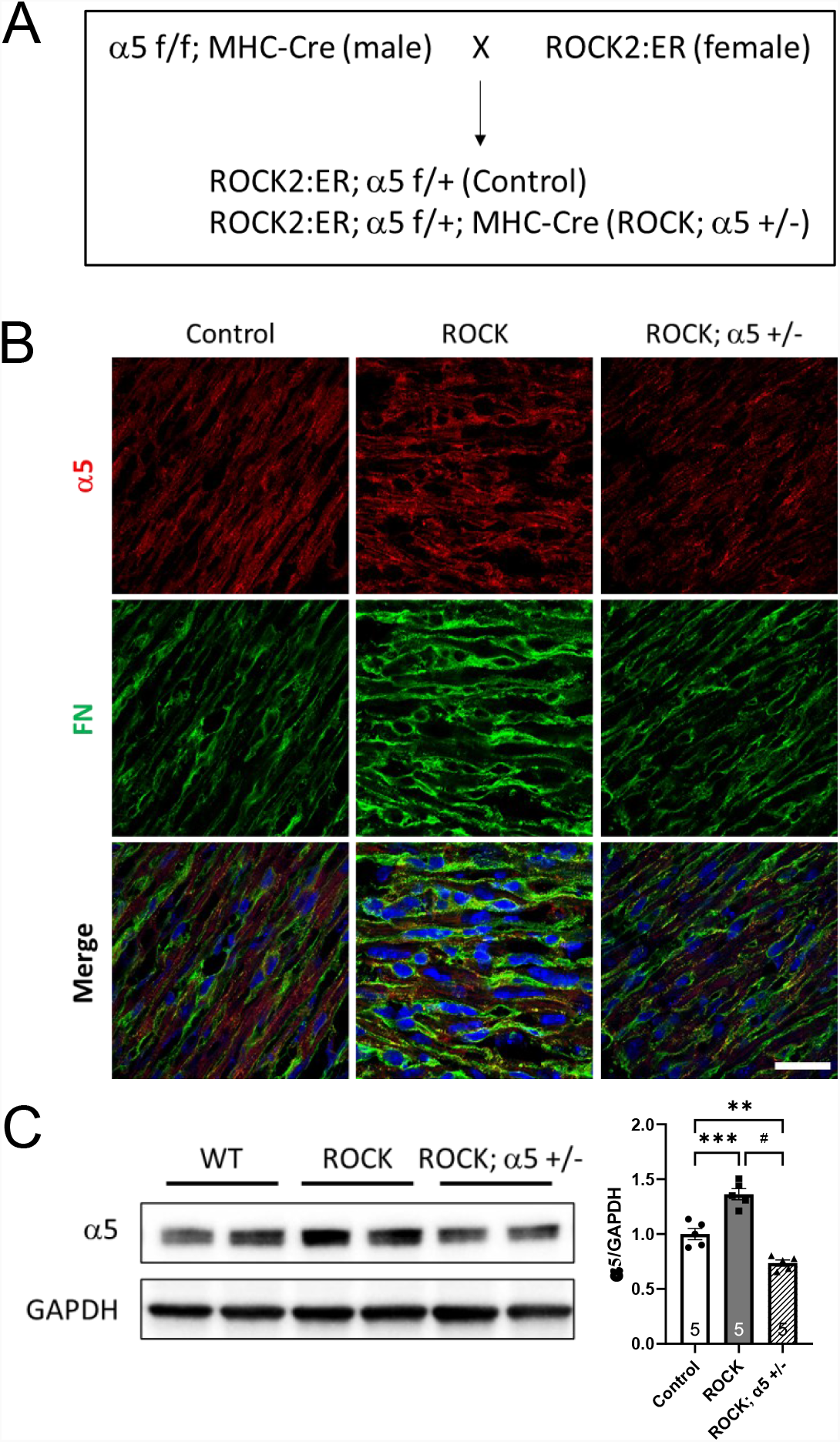
Cardiomyocyte α5 integrin is required for increased fibronectin matrix assembly in ROCK2:ER hearts. (A) Breeding scheme to generate ROCK2:ER; *Itga5 f/+*; MHC-Cre mice (ROCK; α5 +/-). ROCK2:ER; *Itga5 f/+* without MHC-Cre serves as control. (B) Representative immunofluorescence images of heart sections from control, ROCK, and ROCK; α5 +/-P7 mice co-stained with α5 integrin (red) and fibronectin (green). (C) Western blot and quantitative analysis of α5 integrin expression in heart tissue lysate from control (n=5), ROCK (n=5), and ROCK; α5 +/-(n=5) P7 mice. **, P<0.01, ***, P<0.001, #, P<0.0001 by one-way ANOVA with Tukey’s multiple comparisons. Error bars represent S.E.M.. Scale bar: 25μm.

**Figure. 6.**
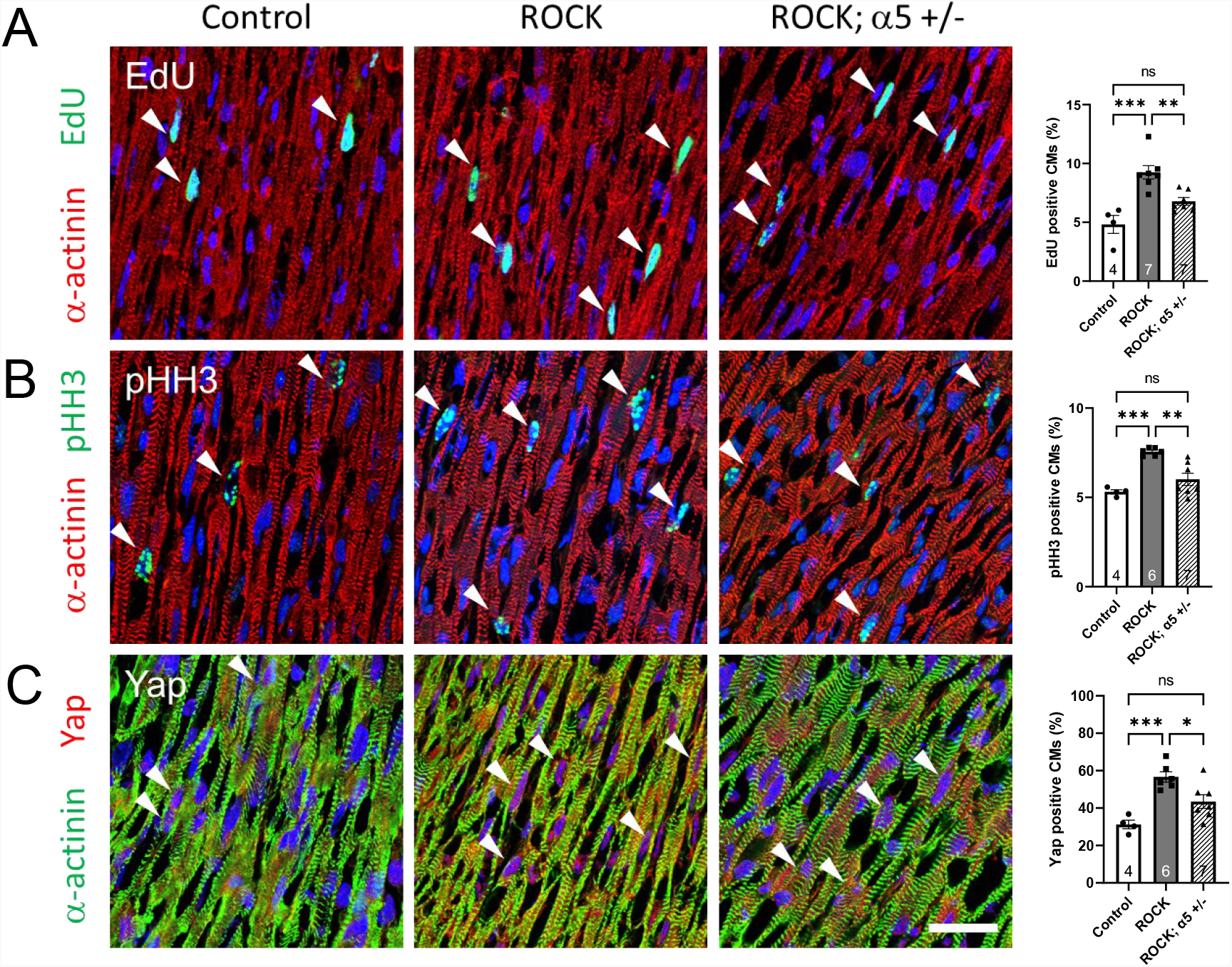
Cardiomyocyte-fibronectin interactions are required for enhanced proliferation in ROCK2:ER mice hearts. Representative immunofluorescence images and quantification of heart sections from control, ROCK and ROCK; α5 +/-P7 mice (n=4-7) co-stained with (A) EdU (green)/α-actinin (red), (B) pHH3 (green)/α-actinin (red), and (C) Yap (red)/α-actinin (green). Arrowheads indicate EdU, pHH3, or nuclear Yap-positive cardiomyocytes. *, P<0.05, **, P<0.01, ***, P<0.001 by one-way ANOVA with Tukey’s multiple comparisons. Error bars represent S.E.M.. Scale bar: 25μm.

**Figure 7.**
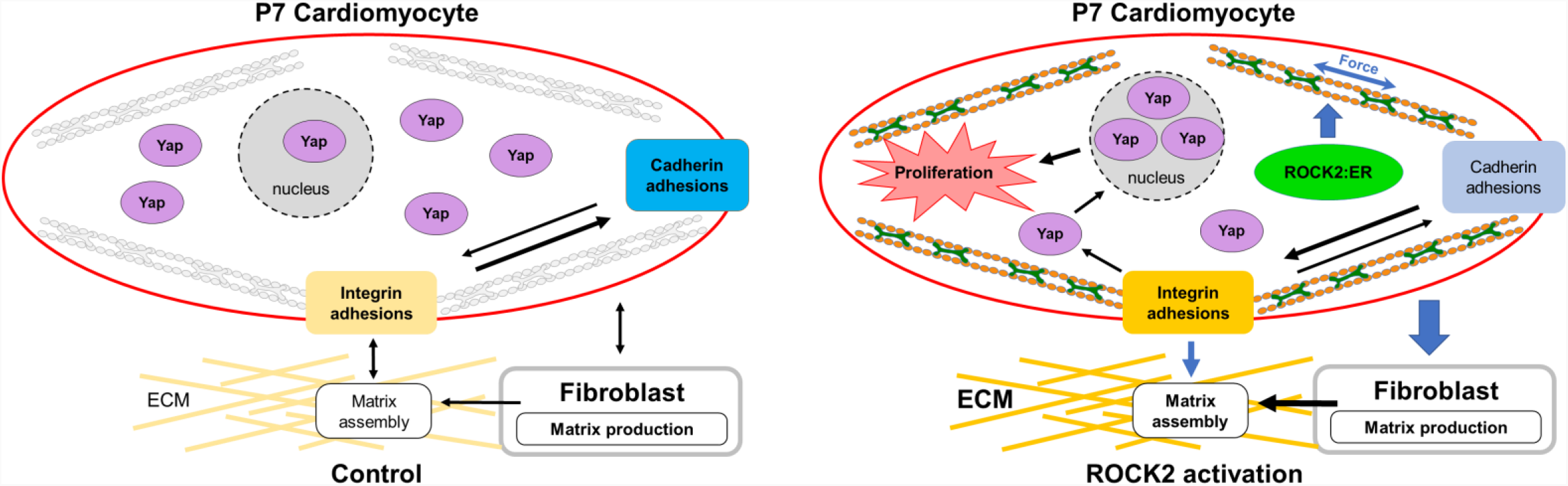
Model for the role of nonmuscle myosin contractility in restricting cardiomyocyte proliferation. Nonmuscle myosin contractility declines as muscle myosin containing myofibrils mature and align with respect to the longitudinal axis of the postnatal cardiomyocyte. In ROCK2:ER hearts, activation of ROCK increases actomyosin contractility which, in turn, drives tissue-level compensatory response that leads to enhanced myocyte-ECM interactions, Yap nuclear translocation, and increased cardiomyocyte cell cycle activity.

## Discussion

Shortly after birth, nonmuscle myosin contractility declines as CMs mature in response to increased cardiac demand and growth during the postnatal period. During the same period, myocardial cell-ECM and cell-cell interactions reorganize resulting in the assembly of strong cell-cell contacts comprised of N-cadherin/catenin adhesion complexes. The postnatal heart thus offers a unique system to investigate how actomyosin generated contractile force regulates cell adhesion, CM maturation, and cell cycle arrest. We previously reported that disruption of N-cadherin junctions via α-catenin depletion was associated with increased actomyosin activity, enhanced Yap nuclear translocation and increased CM proliferation in the postnatal heart (Vite et al., 2018). Based on these findings, it was hypothesized that downregulation of nonmuscle myosin contractility might be required for proper remodeling and maturation of cadherin-based and integrin-based adhesions during postnatal heart development. Here, we utilize a cardiac-restricted ROCK2 conditionally activated GOF murine model (ROCK2:ER) to directly evaluate the consequences of actomyosin contractility on postnatal CM cell adhesions and cell cycle arrest. The tamoxifen activated ROCK2:ER model has been used to study the effects of cellular tension on tissue homeostasis including epidermis, mammary gland, intestine, and pancreas (Boyle et al., 2020; Rath et al., 2017; Samuel et al., 2011; Samuel et al., 2016). To our knowledge, this is the first time the ROCK2:ER model has been used to manipulate actomyosin contractility in a cell type expressing both muscle myosins and nonmuscle myosins.

We report here that activation of ROCK2:ER in postnatal CMs shifts the balance from cadherin adhesions in favor of integrin adhesions, thus promoting the nuclear translocation of Yap, which results in increased CM cell cycle activity. It was recently reported that RhoA activity is required for cyclic stretch-induced Yap nuclear translocation in cultured neonatal CMs (Byun et al., 2019) supporting our *in vivo* findings that activation of ROCK signaling pathway is sufficient to drive Yap to the nucleus. Importantly, in postnatal CMs Yap directly targets genes involved in cytoskeletal remodeling at integrin adhesions (Morikawa et al., 2015). Moreover, Yap loss-of-function in nonmuscle cells leads to downregulation of α5 integrin and reduced assembly of focal adhesions (Nardone et al., 2017). Together, these data suggest that ROCK/integrin/Yap mechanotransduction pathway functions as a positive-feedback loop to maintain cytoskeletal tension via cell-ECM adhesions, thus promoting Yap nuclear translocation.

It is intriguing that nonmuscle myosin activity is re-activated in the adult myocardium following myocardial infarction, where pMLC is found co-localized with vinculin at costameres (Pandey et al., 2018). Whether activation of actomyosin activity in the setting of ischemic injury is compensatory and beneficial remains to be determined. In support of a beneficial effect, expression of modest levels of constitutively active RhoA protected mice from ischemia/reperfusion (I/R) injury, as determined by reduced infarct size and improved contractile function (Xiang et al., 2011). In the same study, RhoA loss-of-function resulted in worse cardiac outcome following I/R injury. Thus, we can speculate that nonmuscle myosin may reinforce cell-matrix interactions at costameres, thus strengthening cardiac contractility following ischemic injury. Future experiments will be necessary to address this and other alternative possibilities.

In mature CMs, actomyosin activity is localized at the ICD along with N-cadherin, suggesting a role in maintaining the structural integrity of the heart. In support of this idea, cardiac-specific loss of NM IIB leads to aberrant N-cadherin expression, altered ICD structures, and cardiac hypertrophy in adult mice at 10 months of age (Ma et al., 2009). In future studies, it will be of interest to explore the physiological consequences of enhanced nonmuscle myosin contractility on cardiac function under normal and pathological conditions.

During the postnatal period, cardiac ECM undergoes significant remodeling resulting in increased tissue stiffness (Hortells et al., 2019). Fibronectin (FN) has emerged as a critical regulator of cardiac development and myocardial growth control (Ieda et al., 2009; Mittal et al., 2013; Trinh and Stainier, 2004; Wu et al., 2020). FN matrix assembly is mediated by integrin receptors and regulated by intracellular signals, cytoskeletal organization, and availability of FN (Singh et al., 2010). FN protein levels decline one week after birth, as collagen levels increase and remain high in the adult myocardium. Following ischemic injury in humans, FN is re-expressed in the infarct and border zone (Willems et al., 1996). Cardiac integrin α5/β1 is the primary receptor for FN. Here we show that cardiac-specific knockdown of α5 integrin (i.e., *Itga5 f/+*) in ROCK hearts restores CM nuclear Yap and CM cell cycle activity to control levels, demonstrating that CM-FN interactions are required for the ROCK proliferation phenotype. In support of our results, co-culture experiments showed that secretion of FN by embryonic cardiac fibroblasts can induce embryonic CM proliferation which is dependent on β1 integrin signaling (Ieda et al., 2009). However, Yap activity was not investigated in this earlier study.

Cardiac fibroblasts (cFbs) are the major cell type contributing to the synthesis of the ECM, which surrounds the CMs and acts as a scaffold, maintaining the myocardial tissue architecture and contractile force of the heart (Hortells et al., 2019; Tallquist, 2020). Periostin (Postn) is a marker of activated cFbs, which have important roles in tissue regeneration and wound healing (Kanisicak et al., 2016). Recent lineage tracing studies identified a heterogenous population of cFbs in the postnatal heart consisting of Tcf21-positive(+) cFbs and Postn+ cFbs (Hortells et al., 2020). While less abundant than Tcf21+ cFbs, Postn+ cFbs are highly proliferative and, importantly, their ablation during the first week leads to decrease CM mitotic activity and a reduction in binucleation. The correlation between high nonmuscle myosin activity and appearance of Postn+ cFbs in both the postnatal heart and in the ischemic adult heart suggest that enhanced CM intracellular tension may provide a mechanical cue to activate cFbs. Further studies are necessary to determine whether activation of actomyosin contractility is sufficient to induce Postn+ cFbs in the heart.

Mechanosensitive proteins such as α-catenin and vinculin undergo force-induced conformational changes resulting in distinct biochemical functions, allowing cells to sense and respond to mechanical stimuli (Charras and Yap, 2018; Pannekoek et al., 2019). Although cytoskeletal tension is important for the formation of both cell–cell and cell–matrix adhesions, excessive tension might induce changes in protein conformation that lead to junctional instability (Weber et al., 2011). In the case of vinculin, site-specific tyrosine phosphorylation distinguishes its mechanical role in cadherin adhesions and integrin adhesions (Bays and DeMali, 2017).

Vinculin conformational activation at cadherin-containing adherens junctions requires tension and phosphorylation at tyrosine residue Y822, which is regulated by the nonreceptor tyrosine kinase c-Abl (Bays et al., 2014; Bertocchi et al., 2017). Here we show that ROCK activation leads to increased VCL Y822 phosphorylation consistent with its role in force-mediated junction maturation in epithelial cells (Bays et al., 2014). Conversely, inhibition of ROCK blocks VCL phosphorylation in epithelial cells exposed to force (Bays et al., 2014). A key avenue for future investigation will be to explore the consequences of blocking vinculin phosphorylation during N-cadherin junction remodeling in the postnatal heart.

In summary, our findings show that nonmuscle myosin contractile forces regulate the balance between myocardial cell-cell and cell-ECM adhesions which, in turn, control Yap nuclear translocation and CM cell cycle activity in the postnatal heart. The ability to induce as well as reverse ROCK activity specifically in CMs will prove invaluable in understanding the role of cellular tension in cardiac development and regeneration, and thus provide critical insight into the potential use of actomyosin agonists for stimulating myocardial proliferation following ischemic injury.

## Materials and Methods

### Generation of cardiac-restricted ROCK2:ER mouse model

*Lox-Stop-Lox ROCK2:ER* mice (Samuel et al., 2016), designated *ROCK2:ER* mice, were bred with *α-Myosin heavy chain (MHC)-Cre* mice (Agah et al., 1997) to generate a cardiac-restricted, conditionally-active ROCK model. *α5 integrin* (*Itga5*) floxed mice (van der Flier et al., 2010) were bred with *MHC-Cre* mice, and resulting *Itga5 f/f*; *MHC-Cre* mice were bred with *ROCK2:ER* mice to generate *ROCK2:ER; Itga5 f/+; MHC-Cre* (ROCK; α5 +/-) mice. The mice were maintained on C57BL/6J genetic background. To conditionally activate ROCK2 kinase in the postnatal heart, P5, P6, and P7 mice were injected intraperitoneally once a day with tamoxifen (Sigma T5648) at 80 mg/kg body weight. To ensure inclusion of both sexes for these studies, the sex of the pups was determined by the presence or absence of the Y-chromosome via PCR. All animal studies were performed in accordance with the guidelines of the IACUC of Rhode Island Hospital.

### Immunofluorescence

Heart tissues were harvested 4 hours after the third and final tamoxifen injection and embedded in OCT (Sakura Finetek). OCT sections (6 μm) were fixed in 4% paraformaldehyde (PFA) for 10min and then washed by PBS 3 times. The heart sections were then permeabilized in 0.1% Triton X-100 (in PBS) for 10min at room temperature. Sections were blocked by T-BSA (5% BSA in 0.01% Triton X-100 in PBS) for 1 hour at room temperature and incubated overnight at 4°C with primary antibodies: phospho-Histone H3 (pHH3) (06-570, Millipore), YAP (4912, Cell Signaling), α-actinin (A7732, Sigma), N-cadherin (610921, BD Bioscience), phospho-MLC2 (3674, Cell Signaling), myosin IIb (M7939, Sigma), α5 integrin (553319, BD bioscience), fibronectin (PA1-23693, Thermofisher), phospho-VCL-Y822 (pVCL) (ab61071, Abcam), VCL (V4505, Sigma). After washing in PBS, sections were incubated with the secondary antibody (Alexa Fluor 488 goat anti-rabbit or 488 goat anti-mouse IgG, Alexa Fluor 555 goat anti-mouse IgG, Alexa Fluor 586 donkey anti-rat IgG, Life Technologies) for 1 hour at room temperature. Finally, the tissue sections were washed in PBS and mounted with ProLong Gold Antifade Reagent containing DAPI (Life Technologies). Images were acquired using a Nikon A1R Confocal Microscope System.

### Monitoring DNA synthesis with EdU Labeling

To assess cardiomyocyte proliferation in heart tissue, mice were injected intraperitoneally with 5-ethynyl-2’-deoxyuridine (EdU, 20 mg/kg body weight) at P7. Four hours after the injection, the hearts were removed, washed in PBS and embedded in OCT. DNA synthesis was revealed with Click-iT EdU Alexa Fluor 488 Imaging Kit (C10337, Life Technologies, Waltham, MA) following manufacturer instructions and then co-stained with cardiac marker α-actinin (A7732, Sigma). A minimum of 10 fields at 40x with a total number of more than 4000 cardiomyocytes per animal were analyzed.

### Western blot analysis

The harvested mice heart tissues were homogenized in a modified RIPA buffer (50mM Tris-HCl pH 7.5, 150mM NaCl, 1mM EDTA pH 8.0, 1% NP-40, 0.5% Na deoxycholate, 0.1% SDS) or SDS-Urea buffer (for the detection MLC2 and phospho-MLC2; 1% SDS, 8mM Urea, 10mM Tris pH 7.5, 140mM NaCl, 5mM EDTA, 2mM EGTA), containing protease inhibitor and phosphatase inhibitor cocktails II and III (Roche Diagnostics). After rotating at 4°C for 2 hours, the mixtures were centrifuged at 13000g for 15 min at 4°C. Primary antibodies used were: phospho-MLC2 (3674, Cell Signaling), MLC2 (3672, Cell Signaling), myosin IIb (M7939, Sigma), ROCK2 (HPA007459, Sigma atlas antibody), phospho-MYPT1 (ABS45, millipore), MYPT1 (612164, BD biosciences), α5 integrin (sc-166681, Santa Cruz), phospho-VCL-Y822 (pVCL) (ab200825, Abcam), VCL (V4505, Sigma), N-cadherin (610921, BD Bioscience), GAPDH (6c5, RDI). For normalization of signals, blots were also analyzed with anti-GAPDH antibody followed by IRDye 680 or IRDye 800CW conjugated secondary antibody (LI-COR, Lincoln, NE). anti-mouse-HRP or anti-rabbit-HRP secondary antibody (Bio-Rad) were also used and developed by SuperSignal West Pico PLUS Chemiluminescent Substrate (Thermo Scientific, Rockford, IL). Membranes were imaged with Odyssey Infrared Imaging System (LI-COR) or ChemiDoc Imaging System (Bio-Rad, Hercules, CA). Quantification was performed with FIJI ImageJ 1.53 software.

### Quantification of cardiomyocyte cell cycle activity and Yap nuclear translocation

For quantification of EdU/pHH3/nuclear Yap positive CMs in IF images, a minimum of 10 fields (40X)/ heart were analyzed using programmed analysis functions in NIKON NIS-Element AR software (Version 5.21.03). Alpha-actinin served as a CMs marker. DAPI indicated nucleus. CM with Yap overlap with DAPI was counted as nuclear Yap positive CM. CM with pHH3/EdU overlap with DAPI was counted as pHH3-/EdU-positive CM. Total number of DAPI within α-actinin-positive cells was regarded as total number of CMs. Ratio of nuclear Yap-, pHH3-, and EdU-positive CMs to total CMs were calculated and plotted in GraphPad Prism 9.

### Statistical analysis

Statistical differences were assessed by unpaired, two-tailed Student’s t test or one-way ANOVA followed by Tukey’s multiple comparison of individual means. A *p* value of < 0.05 was considered statistically significant.

## Figure legends

**Supplemental Figure 1.**
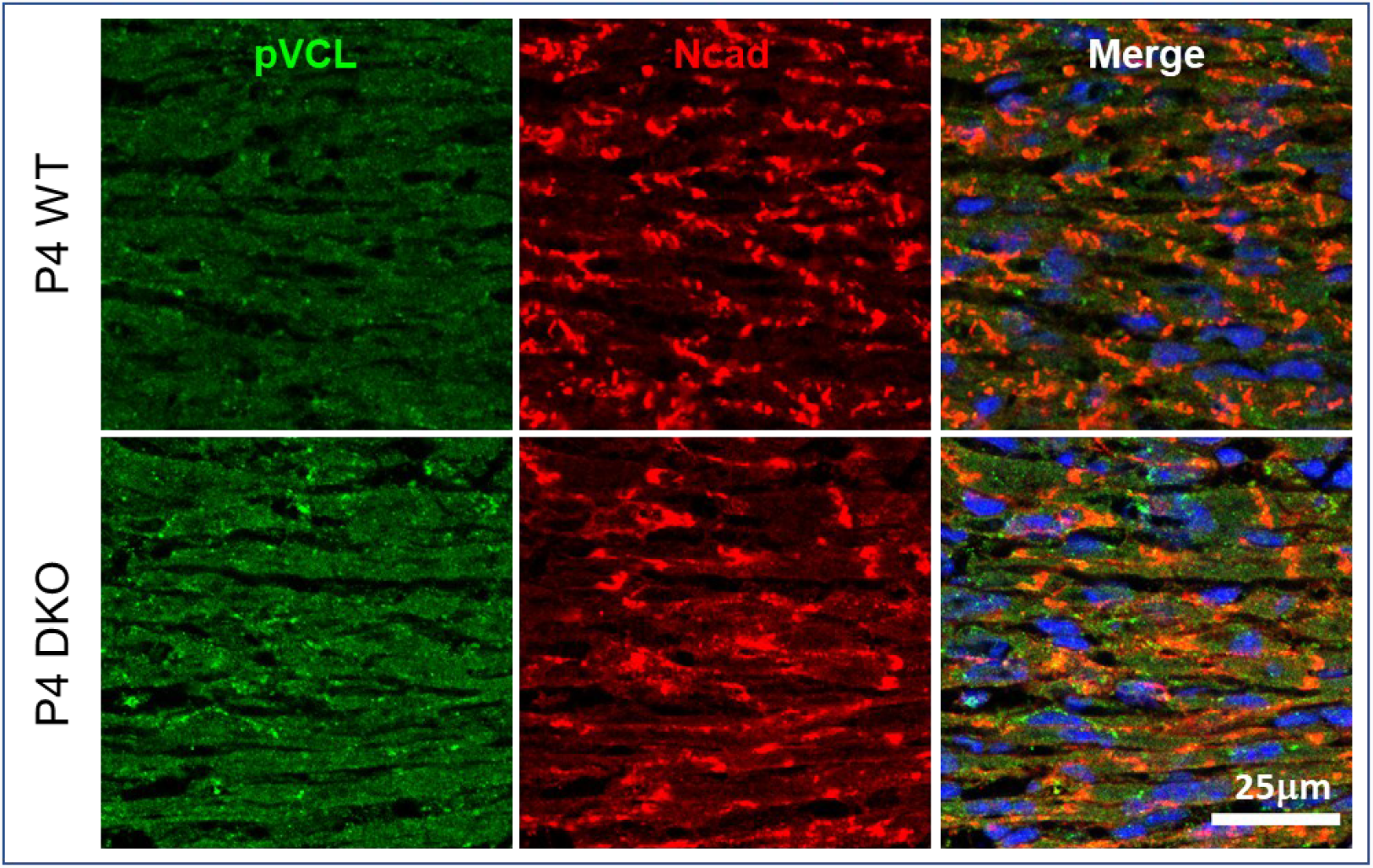
Increased VCL Y822 phosphorylation correlates with aberrant N-cadherin expression in αE-/αT-catenin double knockout (DKO) hearts. Representative immunofluorescence images of heart sections from control and αE-/αT-catenin DKO P4 mice co-stained for pVCL-Y822 (green) and N-cadherin (red). Similar to ROCK-activated hearts, pVCL appeared primarily diffuse throughout the cytosol.

## Acknowledgements

We thank Richard Hynes for the floxed *Itga5* mice provided by Sophie Astrof.

